# Revealing Gene Expression Heterogeneity in a Clonal Population of *Tetrahymena thermophila* through Single-Cell RNA Sequencing

**DOI:** 10.1101/2023.08.06.551249

**Authors:** Hiroki Kojima, Akiko Kashiwagi, Takashi Ikegami

## Abstract

We performed single-cell RNA sequencing (scRNA-seq) on a population of 5,000 *Tetrahymena thermophila*, using the 10x Genomics 3’ gene expression analysis, to investigate gene expression variability within this clonal population. Initially, we estimated the 3’-untranslated regions (3’ UTRs), which were absent in existing annotation files but are crucial for the 10x Genomics 3’ gene expression analysis, using the peaks2utr method. This allowed us to create a modified annotation file, which was then utilized in our scRNA-seq analysis. Our analysis revealed significant gene expression variability within the population, even after removing the effect of cell phase-related features. This variability predominantly appeared in six distinct clusters. Through gene ontology and KEGG pathway enrichment analyses, we identified that these were primarily associated with ribosomal proteins, proteins specific to mitochondria, proteins involved in peroxisome-specific carbon metabolism, cytoskeletal proteins, motor proteins, and immobilized antigens.

## Introduction

Gene expression variability is a widely observed phenomenon in both prokaryotes and eukaryotes (Raj and van Oudenaarden, 2008; Raser and O’Shea, 2005). Single-cell RNA sequencing (scRNA-seq) provides a powerful tool to dissect the mechanisms behind this variability (Saliba et al., 2014; Buettner et al., 2015; Kuchina et al., 2021). While mammalian cells have been extensively analyzed using scRNA-seq, studies on ciliates, particularly those involving sequencing thousands of cells, have been comparatively rare, despite previous studies employing bulk RNA-seq or scRNA-seq with approximately 550 single cells of ciliates (Xiong et al., 2012; Kolisko et al., 2014; Yan et al., 2019; Yang et al., 2023; Grujčić et al., 2024). One reason for this disparity is the larger size of ciliate cells (typically ∼500 μm in length) compared to mammalian cells. Most methods that enable scRNA-seq of thousands of cells are droplet-based, which imposes limitations on the size of cells that can be assessed (∼40 μm) (Ziegenhain et al., 2017).

In this study, we present the results of a scRNA-seq analysis of a ciliate, *Tetrahymena thermophila*, conducted to investigate the gene expression variability. We chose *T. thermophila* as our subject not only because it serves as a primary ciliate model organism for molecular biology and genetics (Turkewitz et al., 2002), but also due to its several advantageous characteristics as follows. Firstly, *T. thermophila* is relatively small, with dimensions of approximately 40–60 μm in length and 20–30 μm in width, making it suitable for droplet-based scRNA-seq techniques. Second, *T. thermophila* mRNA is transcribed from the macronucleus (MAC), and the complete genome sequences of the MAC have already been reported (Eisen et al., 2006; Wang et al., 2021). Last, *T. thermophila* has been observed to exhibit the individual variability in behavior (Jordan et al., 2013). Our preliminary experiments also showed that *T. thermophila* exhibits phenotypic plasticity, with these phenotypes even being inherited by offspring and we were interested in whether the phenotypic variance can be measured by scRNA-seq analysis (to be submitted).

One obstacle to conducting scRNA-seq analysis on *T. thermophila* is the absence of the 3’ untranslated region (3’UTR) annotations, crucial for the 10x Genomics 3’ gene expression analysis method we adopted, which requires accurate 3’ UTR annotation (Zhang et al., 2019). To tackle this, we newly annotate the 3’ UTR by applying the peaks2utr method (Haese-Hill et al., 2023) to the sequence data we obtained. Using this modified annotation, we then conducted scRNA-seq to probe the heterogeneity of the *T. thermophila* population. A primary contributing factor for the expression variability obtained from the scRNA-seq has been proposed to be cell cycle phase. To remove the variations from cell cycle phases and reveal heterogeneity from other sources, we regressed out the contribution from cell cycle phase by utilizing a recently reported set of genes with cell-cycle-dependent expression patterns (Zhang et al., 2023).

In summary, we employed the 10x Genomics system to conduct scRNA-seq on approximately 5,000 *T. thermophila* cells. To isolate the impact of cell-cycle-dependent genes, we conducted regression analysis to mitigate their effects on the gene expression data. Subsequently, we employed a clustering algorithm to analyze the adjusted data. These clusters and gene ontology (GO) and KEGG pathway enrichment analysis revealed the underlying transcriptional variability at the single-cell level in the *T. thermophila* population.

## MATERIALS AND METHODS

### Strain, medium and culture for scRNA-seq libraries preparation

The *T. thermophila* SB210-E strain was used from the Tetrahymena Stock Center (Cornell University, USA). Cells were cultured in the modified Neff medium and PPY medium (Cassidy-Hanley, 2012), which, after storage in liquid nitrogen, were revived in modified Neff medium at 30°C. They were passaged twice in PPY medium at 30°C.

Cells in the late exponential growth phase were loaded onto a 10x Chromium platform using the 10x Single Cell 3’ v3 chemistry, targeting 5,000 cells. The filtration was performed using a 40-μm mini cell strainer (Biomedical Science, Tokyo, Japan) to prevent clogging in the sample line. Our preliminary experiments confirmed that this filtration step did not reduce cell numbers. The filtrated cells were loaded onto a 10x Chromium platform. Libraries were sequenced with NextSeq2000 (Illumina, San Diego, CA) (read one, 28 bp; and read two, 91 bp) to achieve 112,594 mean reads per cell by the Kazusa DNA Research Institute (Chiba, Japan).

### Mapping to genes by Cell Ranger

Sequencing data were analyzed using the 10x Cell Ranger pipeline, version 7.0.1. In order to select the reference MAC sequence, we compared JCVI-TTA1-2.2 (National Center for Biotechnology Information [NCBI] USA) and Tetrahymena_thermophila_mac_genome_v5 (Tetrahymena Genome Database [TGD]) (Stover et al., 2006, Sheng et al., 2020). The percentage of reads confidently mapped to the genome was 81.5% and 14.4% for NCBI and TGD sequences, respectively (Table S1). The low percentage for the TGD sequence indicated that it did not cover the entire genome; hence, we used the NCBI database sequence for further analysis.

For 3’ UTR annotation, we applied the peaks2utr method (Haese-Hill et al., 2023) to the BAM files generated from the MAC sequence by Cell Ranger. Then, we used the modified annotation files to map reads to the MAC sequence and annotated genes by using Cell Ranger once again. The number of cells was set to 5,000, and intron mode was used. Four annotated genes, TTHERM_02141639, TTHERM_02641280, TTHERM_02653301, and TTHERM_002141639 in JCVI-TTA1-2.2, were excluded from subsequent analyses because these sequences were homologous to the ribosomal RNA (rRNA) genes.

### Analysis of the RNA expression matrix

The expression matrix obtained from Cell Ranger was processed using Seurat (Hao et al., 2021). First, we normalized the data by applying sctransform (Hafemeister and Satija, 2019). After normalization, the principal component analysis (PCA) was applied to the 3,000 selected variable features. Then we clustered the data by using the first 10 principal components via the Louvain algorithm (resolution = 0.25). For the visualization, we used UMAP and the markers of each cluster were identified by Wilcoxon Rank Sum test with the Bonferroni correction using Seurat’s findmarker function with min.pct = 0.25 and logfc.threshold = 0.25.

### Cell cycle scoring and regressing out

Cell phases were scored using the CellCycleScoring function (Tirosh et al. 2016) in Seurat. The markers of each cell phase (Table S2) are based on the list from Zhang et al. (2023). This score was then used to regress out the effect of the cell phase. This process was conducted by applying sctransform to the data with setting vars.to.regress to these scores. The same clustering and marker extraction were then applied to the regressed-out data. To check the validity of the resulting cluster, we applied the significance test for clustering, sc-SHC (Grabski et al., 2023). We chose family-wise error as α = 0.25, which is used for the real data application in Grabski et al.

### Gene Ontology and KEGG pathway enrichment analysis

Gene Ontology (GO) and KEGG pathway enrichment analyses were conducted using DAVID, version 2021 (DAVID Knowlegebase, v2023q4) (Sherman et al., 2022). P-values were calculated using Fisher’s exact test with Bonferroni correction.

### Data availability

The FASTQ file, analyzed file, and 3’ UTR annotation file can be found in DDBJ with the accession numbers DRR440460, E-GEAD-611, and BR001895-BR002394, respectively.

## RESULTS AND DISCUSSION

### Mapping of reads

First, we annotated 16,265 3’ UTRs for the MAC sequence available in the NCBI database by applying the peaks2utr method. We confirmed that the distribution of the length of 3’ UTR (median length = 214) was comparable to the previous report (Xiong et al., 2012). Subsequently, we mapped the obtained reads to this modified sequence.

The genome mapping rate was high at 81.5%, but the transcriptome mapping to the NCBI database was only 13.4%, with 67.5% classified as intergenic regions by Cell Ranger. To investigate the origin of this apparent low percentage of the transcriptome mapping, we conducted the read mapping with four different annotations, with or without 3’UTR and with or without rRNA, respectively (Table S1). We found that the mapping with 3’UTR and rRNA yielded 78.8% transcriptome mapping, while without 3’UTR and rRNA was 9.6% and without 3’UTR and with rRNA (the original annotation) was 31.6%. Therefore, the low transcriptome mapping percentage was primarily due to most obtained reads corresponding to rRNA genes (and their associated 3’ UTRs). Also, we confirmed that the annotation of 3’UTR enhanced the transcriptome mapping rate from 9.6% to 13.4%. Although only 13.4% of all reads were mapped to the transcriptome, our results covered 21,875 genes, which correspond to 80.7% of the annotated genes (26,996 genes).

In the 10x single cell 3’ v3 chemistry method, mRNA was selectively obtained using polyA capture, but still approximately 65% of all reads were attributed to rRNA. Kolisko et al. (2014) performed RNA-seq on a single cell of *T. thermophila* using the polyA captured method, and also reported that approximately 90% of reads were mapped to rRNA. These high amounts of rRNA contamination may be because *T. thermophila* has a larger amount of rRNA than other organisms. It contains a large amount of rRNA resulting from a large amount of rDNA, about 9,000 copies (Orias et al., 2011). On the other hand, 2.7% of reads remained unmapped. Cell Ranger analyzes only annotated genes as transcriptomes, so this is likely a sequence read that is not currently annotated as a CDS. Reads unmapped by this analysis may include unannotated CDS, non-coding RNAs, alternative splicing RNAs.

### Clustering results

The expression patterns were divided into seven clusters by analyzing the outputs from Cell Ranger using Seurat (Fig.1(a) and (c)). Each cluster was characterized by a set of genes exhibiting high expression within that cluster (Fig.1(c)). Each cluster contained between 9 to 272 significantly upregulated genes (Table S3).

**Figure 1.**
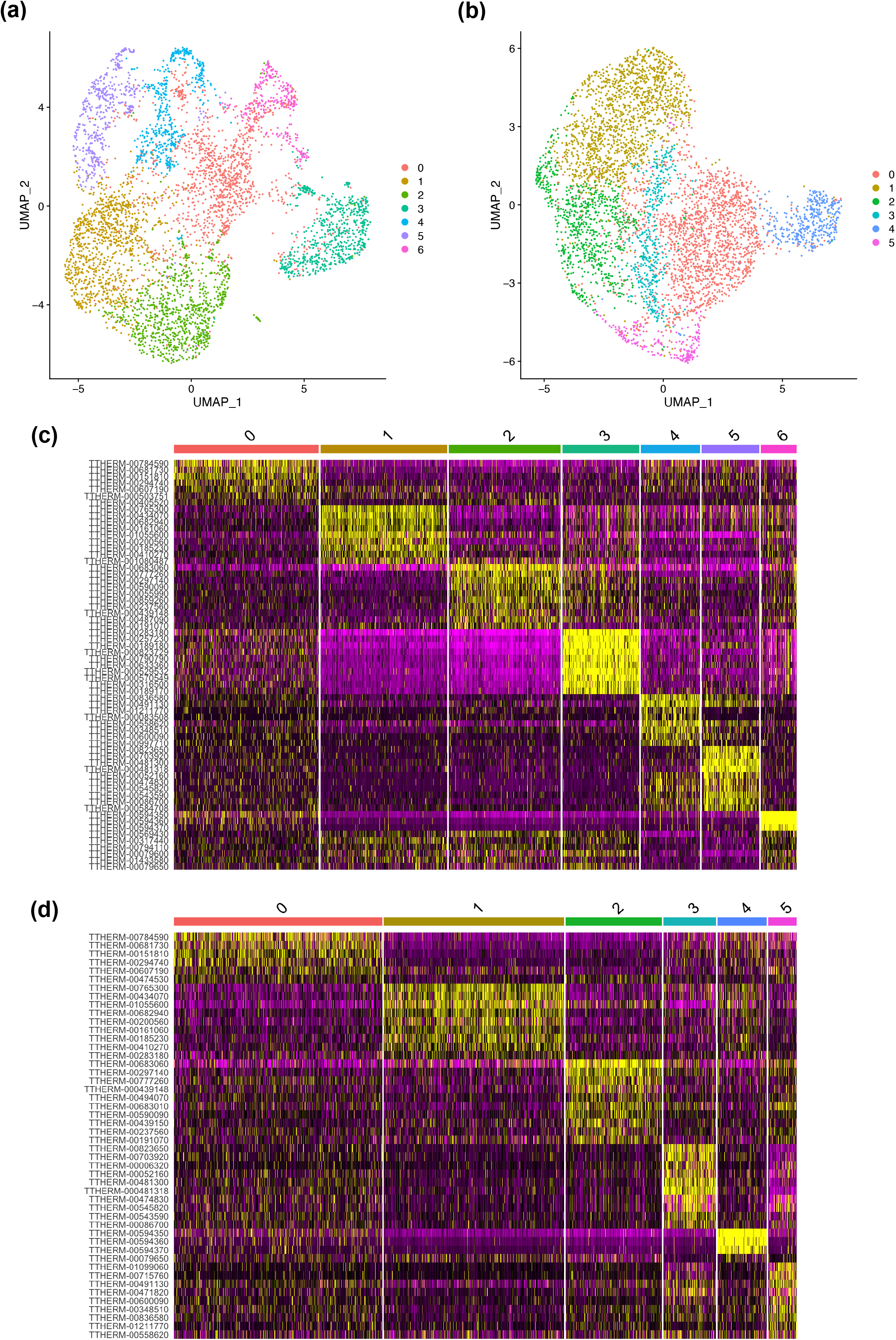
scRNA-seq clustering analysis (a, b) UMAP representation of the RNA expression pattern before and after regression of cell cycle-dependent genes, respectively. (c, d) Heat maps showing differential expression in the population before and after regression of cell cycle dependent genes, respectively.

To characterize each cluster, GO and KEGG pathway enrichment analyses were performed for the six clusters, excluding cluster 0 (Fig. 2 (a)). Based on these analyses, the clusters showed different profiles:

**Figure 2.**
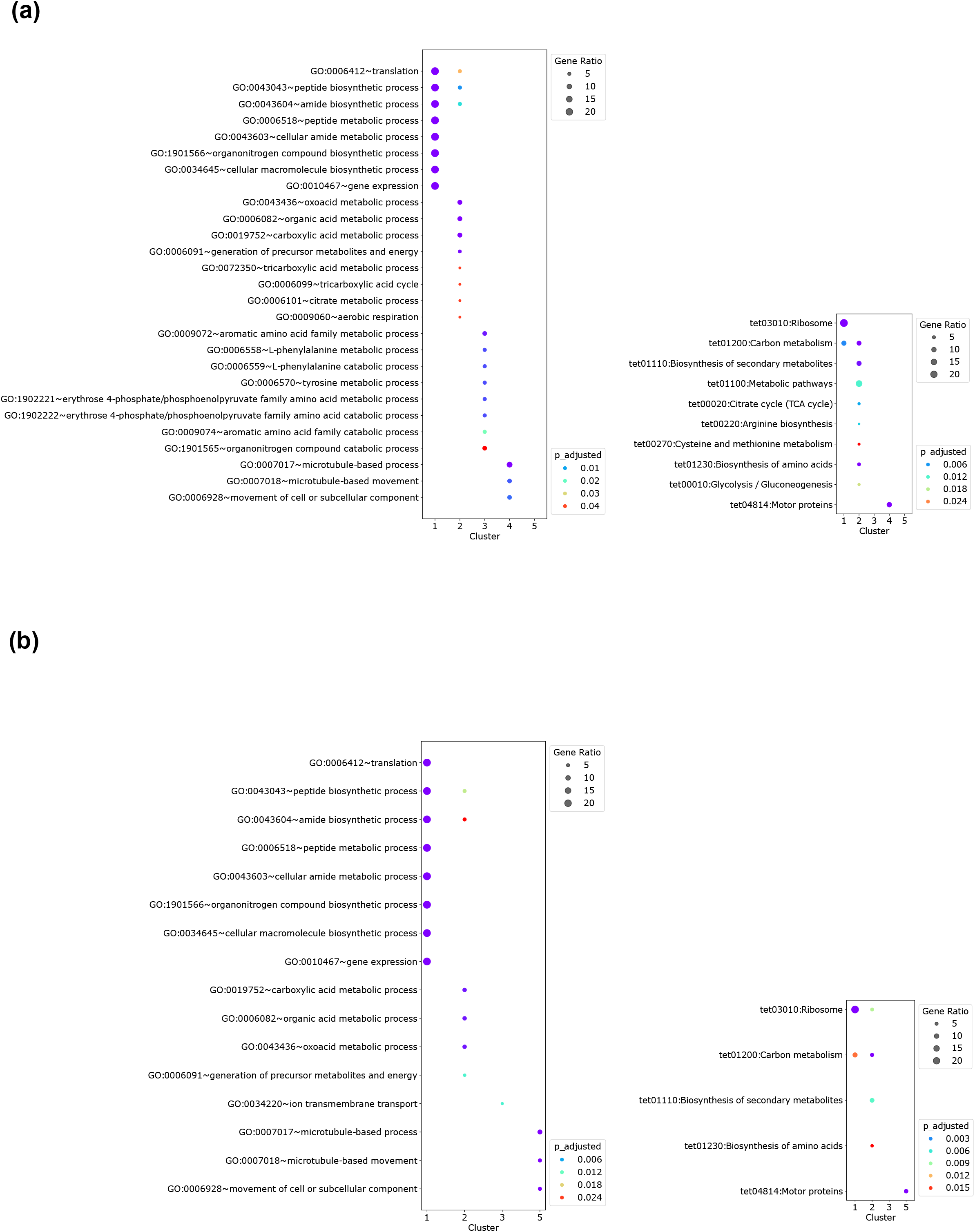
The Gene Ontology (GO) and KEGG pathway enrichment analyses (a) and (b) represent the GO term and KEGG term, before and after regression, respectively. The gene ratios are depicted by differences in circle size, while the adjusted p-values for multiple comparisons using Bonferroni correction are represented by variations in color.

Cluster 0: Functional classification was not appreciable. Among the nine genes analyzed, one was designated as encoding hypothetical proteins, three lacked GO terms and the remaining five were identified as the genes associated with intracellular anatomical structure, translation, protein folding, membrane transport and electron transfer.

Cluster 1: Highly enriched in ribosome and carbon metabolism

Cluster 2: Highly enriched in carbon metabolism including glycolysis/ gluconeogenesis, TCA cycle, and biosynthesis of amino acids

Cluster 3: The enrichment analyses were not applicable well, but characterized by the expression of histones

Cluster 4: Highly enriched in motor proteins

Cluster 5: The enrichment analyses were not applicable well, but characterized by the expression of cytoskeletal proteins unique to ciliates

Cluster 6: The enrichment analyses were not applicable, but characterized by the expression of immobilization antigen and papain family cysteine protease.

To observe gene expression patterns of cells in each cell cycle, we scored cells based on the genes that had been previously reported as cell cycle-dependent (Zhang et al., 2023) and mapped on UMAP (Fig. 3 (a)). Cells expressing genes for MAC-late S phase were found in cluster 3, whereas cells expressing genes for MAC-amitosis phase were found in cluster 4. Cells expressing MAC-G1 phase genes were distributed across various clusters on UMAP (Fig. 3 (a)), indicating that the expression pattern of MAC-G1 phase genes differed among individual cells.

**Figure 3.**
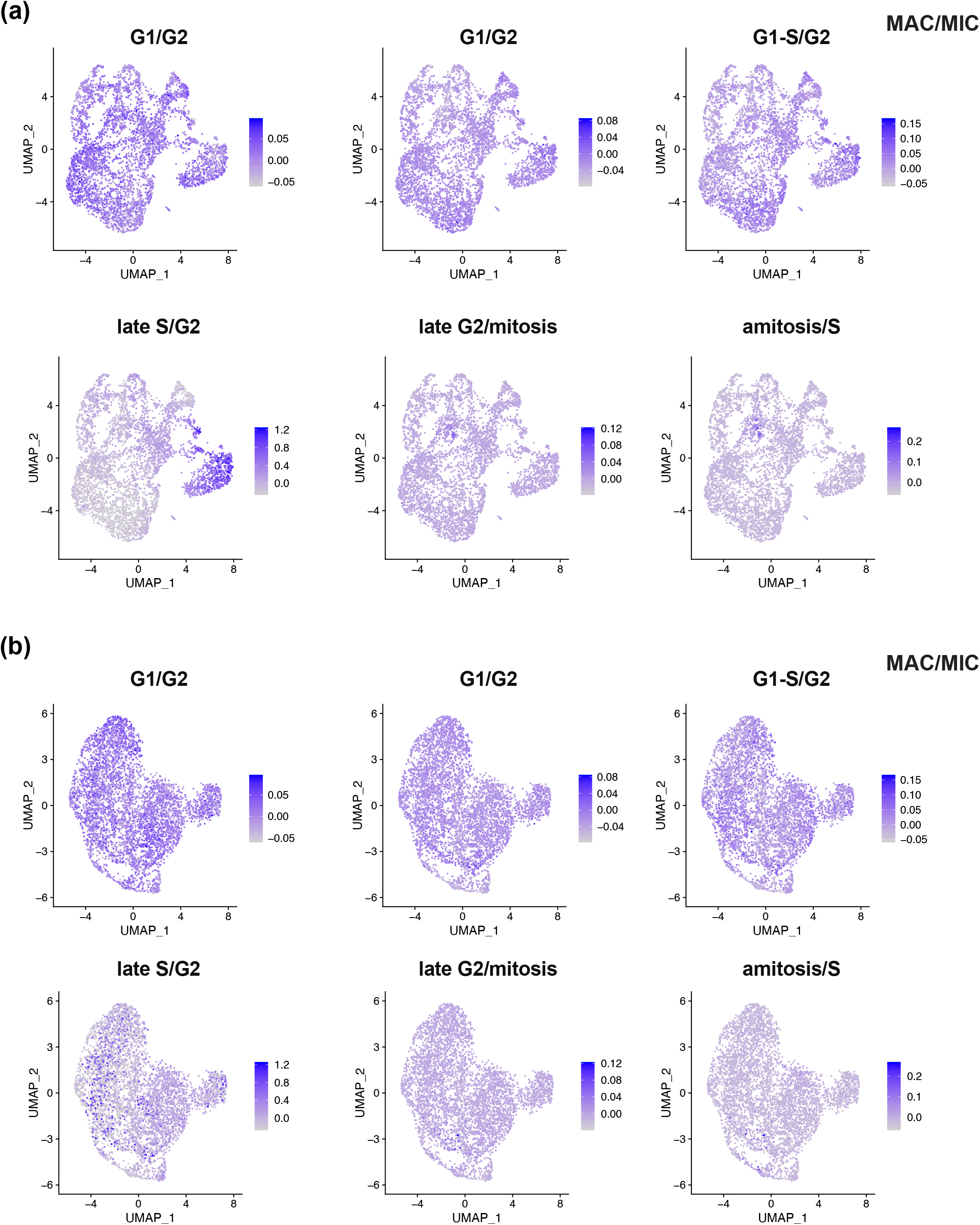
The mapping of genes that have been reported to exhibit cell-cycle dependent expression on UMAP calculated from the gene expression scores of cluster 5B, 7, 5A, 5C, 1 and 2 in Zhang et al., (2023). (a) and (b) represent the gene mapping before and after regression, respectively. The cell-cycle phases are labeled above each panel as MAC/MIC.

When analyzing the top 30 significantly upregulated genes within cluster 3, we found that ten of them were histone genes, including *hhf1, hhf2, hho1, hht1, hht2, hta1, hta2, hta3, htb1*, and *htb2*. Additionally, three genes associated with HMGC high mobility group (HMG) box protein for DNA bending or supercoiling, *hmgb1, hmgb2*, and *nhp6b*, and two genes for structural maintenance of chromosomes, *smc2* and s*mc4*, were identified. Notably, *hhf1, hhf2, hmgb1*, and *hmgb2* are registered as genes expressed during the MAC-S phase in TGD. These findings suggested that cluster 4 probably consisted of cells in the MAC-S phase.

### Regressing out the cell cycle scores

To observe the expression variability without the effects of the cell cycle, we regressed out the cell cycle score using the function provided in Seurat (Hao et al., 2021). After this procedure, mapping of cells that expressed the MAC-late S and MAC-amitosis phase genes on UMAP were scattered across different clusters, indicating that the regression had been successful (Fig. 3 (b)). The resulting expression pattern was divided into six clusters (Fig. 1(b) and (d), Table S4), which were confirmed from the significance test by sc-SHC (α = 0.25).

We conducted GO and KEGG enrichment analyses to each cluster, except for clusters 0 and 4 as it only consisted of six and four genes (Fig. 2(b)). Based on this, the clusters show distinct profiles:

Cluster 0: Functional classification was not appreciable. Among the six genes analyzed, two were designated as encoding hypothetical proteins, two lacked GO terms, while the remaining two were identified as the genes for elongation factor Tu, related to translation, and vps16, a protein localized in the amine-terminal region, which appears to be associated with membrane transport.

Cluster 1: Highly enriched in ribosome and carbon metabolism

Cluster 2: Highly enriched in ribosome, carbon metabolism, and biosynthesis of secondary metabolites and amino acids

Cluster 3: The enrichment analyses were not applicable well, but characterized by the expression of ciliate-specific cytoskeletal proteins including tetrins that located in oral apparatus.

Cluster 4: Characterized by the expression of immobilization antigen and papain family cysteine protease, though the enrichment analyses were not applicable.

Cluster 5: Highly enriched in cilia-specific motor proteins including tubulin and dynein.

Despite both showing expression of ribosomal and carbon metabolite genes, notable differences were observed in the carbon metabolite pathways between clusters 1 and 2. Cluster 1 marker genes included those for certain mitochondrial-specific tricarboxylic acid (TCA) cycle-related genes (citrate synthase I, aconitate hydratase, and succinyl-CoA synthetase) and F_0_F_1_-ATPase/synthase related genes (ATP synthase F_1_, α and β subunits) (Fig. S1). Phosphoenolpyruvate carboxykinase, depicted with a dotted line in cluster 1 of Fig. S1, has been reported to be localized in both the cytosol and the mitochondria (Connett and Blum, 1972). Conversely, cluster 2 marker genes comprised peroxisome-specific genes, such as those involved in the glyoxylate cycle (isocitrate lyase and malate synthase) (Hogg and Kornberg, 1963), and gluconeogenesis-specific (fructose-1,6-bisphosphate). Additionally, both clusters included genes for acetyl-CoA acyltransferase or enoyl-CoA hydratase involved in the β-oxidation of fatty acids.

Given the pivotal roles of mitochondria and peroxisomes, we assume cells in cluster 1 may be in an active metabolite state, primarily producing ATP through the TCA cycle and electron transport system. Conversely, cells in cluster 2 may be in an active metabolite state, focusing on producing acetyl-CoA from fatty acids and synthesizing sugars and proteins. Our speculation rests on the previous findings that the glyoxylate cycle in *Tetrahymena* serves both an auxiliary pathway to the TCA cycle and a crucial role in synthesizing essential substances such as proteins and glycogen, necessary for growth (Raugi et al., 1975, Gotoh et al., 1990). The auxiliary role in the TCA cycle was underscored by the report that, when quantifying carbon flow in *Tetrahymena*, the flow of carbon through the glyoxylate bypass is approximately one-third of the flow through the TCA cycle, and furthermore, the malate generated in the glyoxylate cycle is transported to the mitochondria (Raugi et al., 1975). The role for the substance synthesis was clarified by the report that inhibition of fatty acid β-oxidation in peroxisomes and inhibition of isocitrate lyase activity resulted in a decrease in the supply of acetyl-CoA derived from the β-oxidation in peroxisomes, leading to reduced protein and glycogen synthesis, and consequently, a slower growth rate in *Tetrahymena* (Gotoh et al., 1990).

In summary, our scRNA-seq cluster analysis reveals the presence of approximately six distinct semi-macro states of gene expression during the log phase. This observation challenges the conventional understanding that the internal state of microorganisms remains uniform throughout this growth phase due to a constant growth rate (Madigan et al., 2021). Our findings align with recent scRNA-seq studies on *Bacillus subtilis*, which also report similar gene expression heterogeneity (Kuchina et al., 2021), underscoring a broader pattern of variability within microbial populations that extends beyond individual species.

The fluctuation of gene expression patterns, and thus phenotypic variability, is widely acknowledged across various organisms (Balázsi et al., 2011). However, the underlying causes of this variability—whether arising from inter-individual interactions or inherent instability within individual cells—remain to be fully elucidated.

Malate may serve as a crucial metabolite for inter-individual interactions within cell populations, besides the role in the TCA cycle and glyoxylate cycle. Outside the cell, we have observed the leakage of malate into the culture supernatant (data not shown) and this secreted malate likely plays a role in intercellular communication. In such interactions, cells producing higher levels of malate may secrete it into the surrounding medium, while cells producing lower levels may utilize the excreted malate. Although chemotaxis towards amino acids and peptides is well-studied in *Tetrahymena*, research on chemotaxis towards dicarboxylic acids, including malate, is still forthcoming (Köhidai, 2016). However, in *Pseudomonas fluorescence*, chemotaxis towards dicarboxylic acids including malate, succinate, and fumarate has been observed, suggesting these metabolites function as cell-to-cell signaling molecules (Oku, 2012). Furthermore, studies in experimental molecular evolution using *Escherichia coli* have shown that maintenance of genetic diversity through cell-to-cell interactions mediated by secreted L-glutamine occurs (Kashiwagi et al., 2001). Thus, malate-mediated cell-to-cell interactions could significantly contribute to the dynamics of *Tetrahymena* populations and the emergence of diverse gene expression patterns.

The evolutionary selection of phenotypic fluctuation suggests a profound biological significance. For instance, the partitioning of metabolic reactions into discrete tasks could confer enhanced robustness against environmental perturbations. In our preliminary experiments, we observed that *Tetrahymena* populations exhibit differentiation into entities with varying levels of kinetic energy, which further illustrate how such phenotypic diversity could bolster resilience to environmental changes. Notably, the aggregation of slower-moving individuals could facilitate swarm formation, highlighting a potential adaptive advantage in phenotypic variability.

In this context, our study not only contributes to the growing body of evidence for gene expression heterogeneity among microorganisms but also raises important questions about the adaptive roles of such variability. It suggests that the fluctuating phenotype, far from being a mere byproduct of cellular processes, may be an evolutionarily favored trait that enhances the survival and adaptability of microbial populations. Future research should aim to dissect the mechanisms driving this variability and explore its implications for microbial ecology and evolution. Understanding the balance between genetic stability and phenotypic plasticity could unveil new dimensions of microbial survival strategies in fluctuating environments.

## CONCLUSIONS

First, the use of the peaks2utr enabled us to newly annotate 16,265 3’ UTR for *T*.*thermophila* genome sequences (NCBI). Second, our scRNA-seq analysis revealed that cell phases (especially the MAC-late S phase) were the main contributors to gene expression variability. Third, after removing this cell phase-related variability, resulting heterogeneity was mainly attributed to ribosomal proteins, mitochondria-specific or peroxisome-specific carbon metabolism, cytoskeletal proteins, motor proteins, and immobilized antigens.

## ACKNOWLEDGEMENTS

This study was partially supported by the Uehara Memorial Foundation and by the JSPS Kakenhi Grant-in-Aid for Scientific Research (A) 21H04885. The authors thank Dr. Geoffrey Kapler for providing a list of cell cycle-dependent genes.

## Notes

### Competing Interest Statement

The authors have declared no competing interest.

### Summary of Updates

Enhanced the Results section with revised analyses; Thoroughly revised the language and organization of the text to improve readability; Figure 1 revised; Figure 2,3 added; Supplemental files updated

