## Supplemental Figure 1 for "Revealing Gene Expression Heterogeneity in a Clonal Population of *Tetrahymena thermophila* through Single-Cell RNA Sequencing"

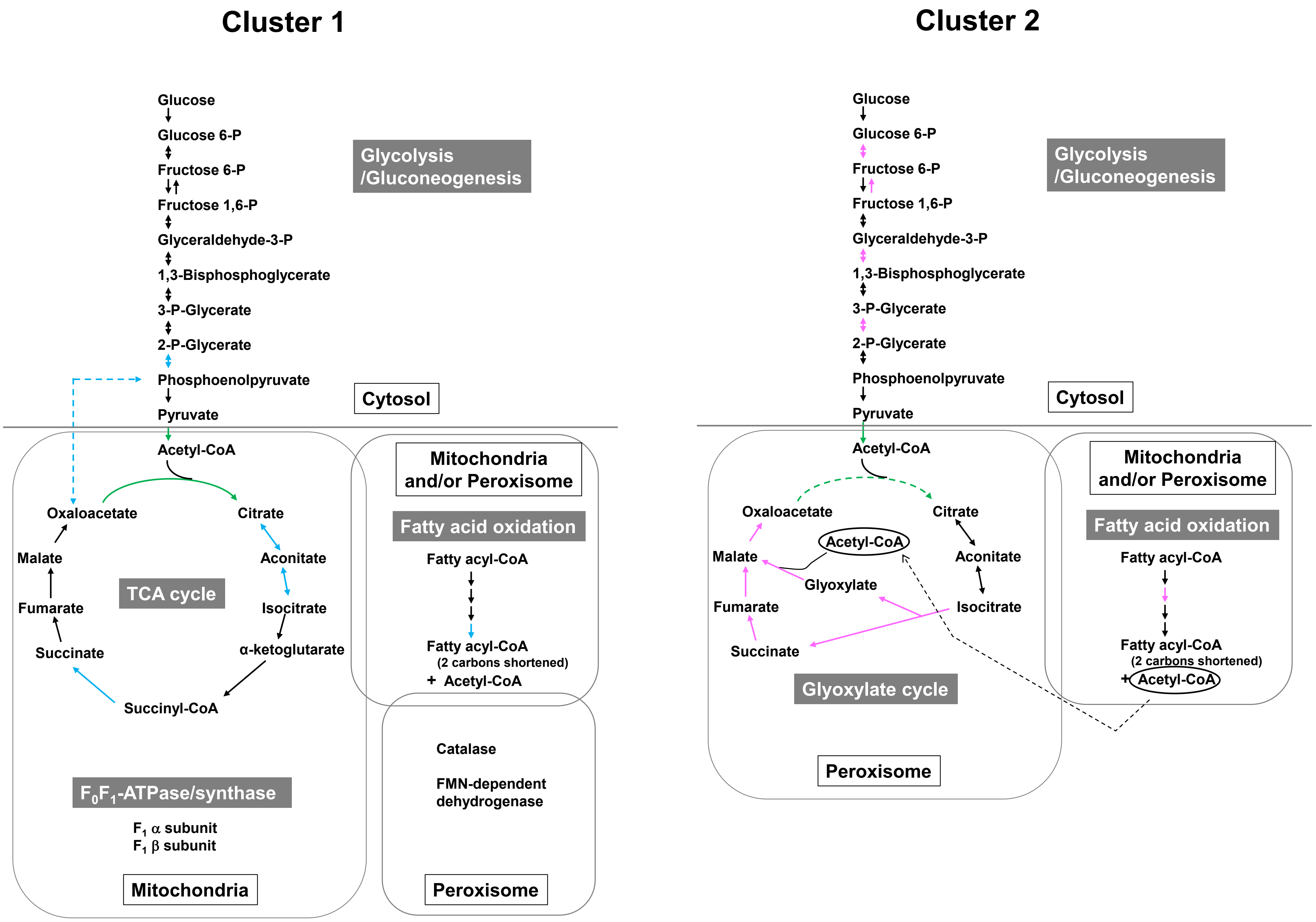


Figure S1. The differences in carbon metabolism pathways between cluster 1 and 2.

The paths of marker genes in each cluster are indicated by blue, green, and pink arrows. Blue and pink arrows represent paths observed exclusively in each cluster, while green arrows signify paths observed in both clusters. Dashed lines indicate pathways for which it is uncertain whether they occur within the mitochondria, peroxisome, or cytosol.
